# Human IFNε: Spaciotemporal expression, hormone regulation and innate immunity in the female reproductive tract

**DOI:** 10.1101/445007

**Authors:** Nollaig M. Bourke, Sharon L. Achilles, Stephanie U. Huang, Helen E. Cumming, Irene Papageorgio, Linden J. Gearing, Suruchi Thakore, Niamh E. Mangan, Sam Mesiano, Paul J. Hertzog

## Abstract

Interferon epsilon (IFNε) plays an important role in regulating protective immunity in the female reproductive tract in mouse models; but the expression and regulation of this IFNε in the human FRT had not yet been characterised. Here we show that IFNε is selectively and highly expressed in the human FRT, a unique characteristic among the many types of IFN. IFNε has distinct expression patterns in upper compared with lower FRT where it is predominantly expressed in the basal layers of the stratified squamous epithelia. We demonstrate direct regulation of IFNε expression is suppressed by progesterone consistent with its inverse correlation with progesterone receptor expression, but only in the endometrium where its expression therefore fluctuates throughout the menstrual cycle. We show that IFNε regulates immunoregulatory IFN regulated genes (IRGs) in FRT epithelial cells. The characterisation of huIFNε expression in both the upper and the lower FRT epithelia and its protective properties make this IFN well placed to be an important player in mediating hormonal control of FRT immune response and susceptibility to FRT infection.

**Summary:** Bourke et al. characterise the novel type I interferon epsilon (IFNε), as the only IFN constitutively expressed throughout the human female reproductive tract (FRT), where it is hormonally regulated and modules IFN dependent FRT immunity.

## Introduction

The female reproductive tract (FRT) mucosa is a unique site of immune regulation as it must be capable of mounting robust responses against pathogenic infections yet maintain tolerance with commensal bacteria, semen and developing pregnancies (Roy and Matzuk, 2011). Epithelial cells that line the vagina, cervix and uterus represent the initial barrier against invading pathogens and are important regulators of immunity through specialised antigen presentation function and secretion of mucins, antimicrobial peptides (AMPs), and chemokines that modulate recruitment and activation of the innate and adaptive immune cells (Wira et al., 2010). The fluctuation of sex hormones in women, primarily estradiol (E2) and progesterone, across the menstrual cycle likely impact immune function in the FRT, although the mechanisms mediating these effects are not comprehensively defined (Wira et al., 2015). During the progesterone-dominated luteal phase of menstrual cycle when the endometrium is prepared for fertilisation and implantation, immune responses are suppressed and a tolerogenic state is established (Wira et al., 2015). However, this environment leaves the FRT susceptible to pathogen invasion. For example, medroxyprogesterone acetate treatment increases Chlamydi*a trachomatis* and HSV2 infection susceptibility in mice (Kaushic et al., 2003; Kaushic et al., 2000). Far less is known about the link between progestins and STI susceptibility in humans, but there are suggestions that some progestin-based contraceptives are linked with increased susceptibility to HIV (Polis et al., 2016; Polis et al., 2014; Ralph et al., 2015). However, the factors that mediate the hormonal effects on host defence are not known. Antiviral and immunoregulatory cytokines, such as type I interferon (IFNs), are prime candidates.

The type I IFNs are a family of cytokines that include the conventional α and β subtypes, as well as the more recently identified IFNε. All type I IFNs bind IFN-α receptor 1 (IFNAR1) and IFNAR2, activate JAK-STAT signalling and regulate the expression of potentially thousands of IFN regulated genes (IRGs) (Schindler et al., 2007). The ‘effector’ proteins induced by IRGs can modulate a wide range of biological responses, including antiviral, cell cycle regulation, survival/apoptosis, immune effector cell activity and chemotaxis. Regarding STIs in the FRT, epithelial cells that line the vaginal, cervical and endometrial mucosa are key sentinels that produce conventional type I IFNs (e.g. α’s and β) upon pathogen challenge (Wira et al., 2010). Now, a “new” and compelling IFN to consider in FRT immunity is IFNε, which we characterised in mouse models to be constitutively expressed in the endometrium and protective against viral and bacterial STIs (Fung et al., 2013). We determined that murine IFNε was not induced by pathogen recognition receptor (PRR) pathways like conventional type I IFNs, but was hormonally regulated and accordingly, IFNε levels fluctuated in the endometrium across the estrus cycle. Furthermore, we and others found that in *in vitro* experiments in human cells, IFNε can block HIV replication at several steps of viral replication by induction of antiviral IRGs (Garcia-Minambres et al., 2017; Tasker et al., 2016). However, the critical question of the location and regulation of IFNε in the FRT and its functions there, remains unreported.

Since the role of IFNε in regulating protective immunity in the human FRT will be dependent on which tissues and cells express it, we undertook a thorough investigation of the expression of type I, II and III IFNs, their regulators and their effectors across the FRT.

## Methods

### Participant recruitment and sampling

We performed a cross-sectional study (ClinicalTrials.gov number: NCT02416154) of healthy, reproductive-aged women with normal menstrual cycles who were free of exogenous hormonal contraceptives. Being free of exogenous steroid hormones and in a defined (follicular vs luteal) phase of menses was central to the study design and therefore laboratory confirmation by ultra-high-performance liquid chromatography tandem mass spectrometry (UPLC/MS/MS) was performed to evaluate serum progesterone, estradiol as well as a panel of synthetic progestins that cover the majority of regionally available contraceptive progestins. The University of Pittsburgh Institutional Review Board and the Monash Health Human Research Ethics Committee both approved this study. All participants were enrolled at the Center for Family Planning Research at Magee-Women’s Hospital in Pittsburgh, Pennsylvania and signed informed consent before study participation.

Between August 2015 and August 2016, 44 participants were assessed for study eligibility and 34 women, age 18-35 years, were enrolled. Of the enrolled participants, 17 were in the follicular phase and 16 were in the luteal phase of menstrual cycle; 1 participant was discontinued after enrolment for a positive screening test for *Chlamydia trachomatis*. Eligible women were healthy, HIV negative, non-pregnant and reported regular menstrual cycles every 21-35 days. Women were excluded if within 30 days of enrolment they: 1) used any hormonal or intrauterine contraceptive methods; 2) underwent any surgical procedure involving the pelvis (including biopsy); 3) were diagnosed with any urogenital tract infection; 4) used any vaginal or systemic antibiotics, oral or vaginal steroids, or any vaginal product or device (including spermicide, microbicide, douche, sex toy, cervical cap, menstrual collection device, diaphragm, or pessary) except tampons and condoms. Women were also excluded if they used depot medroxyprogesterone acetate within 10 months of enrolment, were pregnant or breastfeeding within 60 days of enrolment or had a new sex partner within 90 days of enrolment. Exclusion criteria included having unprotected heterosexual intercourse since last reported menses, having vaginal or anal intercourse within 36 hours prior to the enrolment study visit, having a prior hysterectomy or malignancy of the cervix or uterus, and having any history of immunosuppression, including immunosuppression associated with chronic disease. Medical, gynaecologic and sexual histories were obtained and screening procedures were conducted including urine pregnancy testing; rapid HIV screening (OraQuick®, OraSure Technologies, Bethlehem, PA, USA); collection of genital tract swabs for detection of *Neisseria gonorrhoeae*, *Chlamydia trachomatis* (Hologic Inc., San Diego, CA, USA) and rapid testing for *Trichomonas vaginalis* (OSOM, Sekisui Diagnostics, Lexington, MA, USA). Participants were enrolled on the same day as screening when all eligibility criteria were met including no vaginal bleeding on exam and being in the follicular (day 3-12) or luteal (≤10 days prior to anticipated start of menses) phase of their menstrual cycle by self-report. Given the low risk study population, participants with no clinical signs of genital infection were enrolled with pending screening tests and were discontinued post-enrolment if a screening test rendered them ineligible. Final group allocation to phase of menses was based on UPLC/MS/MS serum hormone analysis with follicular phase serum progesterone <1000pg/mL and luteal phase serum progesterone >2000pg/mL.

Five biopsy samples from the genital tract were obtained: two vaginal biopsies from the upper vagina, two cervical biopsies from the squamocolumnar junction, and one endometrial biopsy. Vaginal and cervical biopsies were obtained with a standard gynaecologic biopsy instrument and each measured approximately 2 x 3 x 2 mm. The endometrial biopsy was obtained using a standard endometrial sampler (Pipelle®, Cooper Surgical, Trunbull, CT, USA). The participants were given the option of a cervical aesthetic injection with 10cc of 1% lidocaine solution prior to the endometrial biopsy. If this was elected, it was administered after the vaginal and cervical biopsies were obtained. One of each of the vaginal and cervical biopsies and ½ of the endometrial biopsy sample were each placed in 1ml RNALater and stored at 4 degrees overnight before being transferred to -80 degrees. The second vaginal and cervical biopsies and the remaining ½ of the endometrial biopsy sample were separately placed into histology cassettes and incubated in 10% formalin fixative for 24 hours at room temperature. The samples were then transferred to 70% EtOH solution at room temperature. Blood samples were also evaluated by UPLC/MS/MS for quantification of estrogens and progestogens as previously described (Achilles et al., 2018). Serum progesterone was also evaluated by the clinical hospital laboratory. Analysis of IFNε and related parameters were carried out in the Hudson Institute of Medical Research in Melbourne, Australia; all analysis was conducted by researchers blinded to cycle stage of participants.

### Cell culture and reagents

We performed a series of in vitro experiments to determine the response of cells from different compartments of the human FRT to correlate their responses with spatial features of their expression. VK2 [vaginal (<u>ATCC^®^ CRL-2616™</u>)], Ect1 [ectocervical (ATCC^®^ CRL-2614™)] and End1 [endocervical (ATCC^®^ CRL-2615™)] were maintained in Keratinocyte-SFM, supplemented with 0.2ng/ml of human rEGF, 20μg/ml of rBPE and 1% Antibiotic-Antimycotic (Gibco, Life Technologies). For stimulation cells were plated at 1.5×10^5^ cells/well in a 12 well plate and stimulated for 3h with 100IU/ml recombinant IFNε (made as described previously (Stifter et al., 2017)) or IFNβ (REBIF, Merck). For primary uterine epithelial cell (UEC) cultures, human endometrium specimens were obtained with approval from the Institutional Human Research Ethics Committee. Endometrial cells were isolated by mincing, digestion with collagenase and DNase I and filtration, as previously described (Fung et al., 2013). UECs were stimulated on day 3 of culture.

### Gene expression analysis

RNA was extracted from biopsy samples from the cross-sectional studies described above, using TRI reagent and the RNeasy Kit (QIAGEN, Melbourne, Australia). cDNA was synthesised following DNase treatment (Promega, USA) using MMLV reverse transcriptase and random hexamers (Promega, USA). *Biomark Fluidigm qPCR:* Gene expression was analysed using the Biomark Fluidigm system. Ct values from the Biomark qRTPCR were calculated using the Biomark Fluidigm Real Time PCR Analysis Software. Comparative analysis was performed using R Studio with the HTQPCR package (Dvinge and Bertone, 2009), including methods for Principle Component Analysis, Visualization as well as Spearman Correlation. Results were normalised to the housekeeping genes HMBS and RPLPO. Unreliable probes were removed as defined by post normalised Ct values >10 or undetected in greater than 30% of samples. *SyBr green qPCR:* For in vitro studies using cell lines, RNA was extracted using an RNeasy Kit and qPCR was performed using SYBR green (ABI, ThermoFisher, Australia) on an Applied Biosystems 7900HT Fast Real-Time PCR Machine and primer sequences are listed in supplemental Table 1. Expression was calculated using the 2−ΔΔCT method using *18S* as endogenous control.

### Immunohistochemistry analysis and quantification

Thin sections (~4μM) were cut for each tissue and adhered onto Superfrost^TM^ Plus glass slides (ThermoFisher, Australia). Sections were dehydrated using a series of 100% xylene, 100% ethanol, 70% ethanol and MilliQ water solutions. Heat induced antigen retrieval was performed using a 10mM Trizma^®^ base (Sigma-Aldrich)/1mM EDTA buffer at pH 9.0. Sections were blocked using CAS-block (Invitrogen), for 1 hour at room temperature, then incubated overnight at 4°C with the following primary antibodies: rabbit anti-human IFNε (Novus Biologicals), used at 0.5μg/mL; mouse anti-human estrogen receptor α (Dako), provided at ready-to-use concentration; and mouse anti-human progesterone receptor (Dako), used at 1.56μg/mL; all diluted in CAS-block. Corresponding isotype controls were rabbit IgG (Vector Laboratories), used at 0.5μg/mL; and mouse IgG1 (Dako), used at 1.56μg/mL; both diluted in CAS-block. Sections were washed with 0.05% Tween-PBS for 15 minutes and incubated with 60μg/mL biotinylated secondary antibodies (anti-rabbit and anti-mouse, Vector Laboratories) for 1 hour at room temperature. Slides were washed in 0.05% Tween-PBS for 10 minutes, then incubated for 45 minutes with VECTASTAIN Elite^®^ ABC-HRP Kit, an avidin-biotinylated peroxidase H complex (Vector Laboratories). Slides were washed again for 10 minutes in 0.05% Tween-PBS, and DAB substrate (Dako) was applied for 30 seconds to initiate precipitate formation/colour development via peroxidase activity. Enzyme activity was stopped in distilled water. Coverslips were applied to slides with DPX mounting media (Merck) and allowed to dry overnight. Slides were manually cleaned, scanned at the Monash Histology Platform using the Aperio Scanscope AT Turbo (Leica Biosystems) and analysed using Aperio ImageScope (v12.3.0.5056, Leica Biosystems) with the Aperio Cytoplasm Algorithm (v2, Leica Biosystems). The epithelium, glands, and stroma were delineated using the pen tool and assessed independently of each other for both nuclear and cytoplasmic staining – an H-score was obtained for each to perform statistical analysis.

### Statistical analysis

For Fluidigm analysis, statistically significant gene changes were determined between conditions using linear modelling, employing the Limma Statistical package [18]. A t-test was used, with a 1.5-fold change cut off and Benjamini-Hochberg correction for false discovery. Gene expression was considered to be significant below a p value <0.05. For all other data, statistical analyses were performed using GraphPad Prism (v5.02, La Jolla, CA, USA). Mann-Whitney U-tests, Wilcoxon matched pairs signed rank test and one-way ANOVA with Bonferroni’s post entry test were used as indicated. Spearman’s rank correlation coefficient analysis was used for all correlation analysis.

## Results

### IFNε is expressed throughout the human FRT and regulated in response to sex hormones exclusively in the endometrium

We obtained matched FRT tissue and blood samples from a cohort of 33 women (17 in the follicular stage of menstrual cycle and 16 in the luteal stage) with strict inclusion and exclusion criteria to minimize potential confounding factors that could modulate FRT environment. The demographic characteristics of the 33 enrolled and eligible participants did not differ by allocated menstrual phase arm and are shown in Table 1. The range of serum progesterone concentrations for women allocated to the follicular group vs the luteal group were 26-973 pg/mL and 2075-16,793 pg/mL respectively.

**Table 1.**
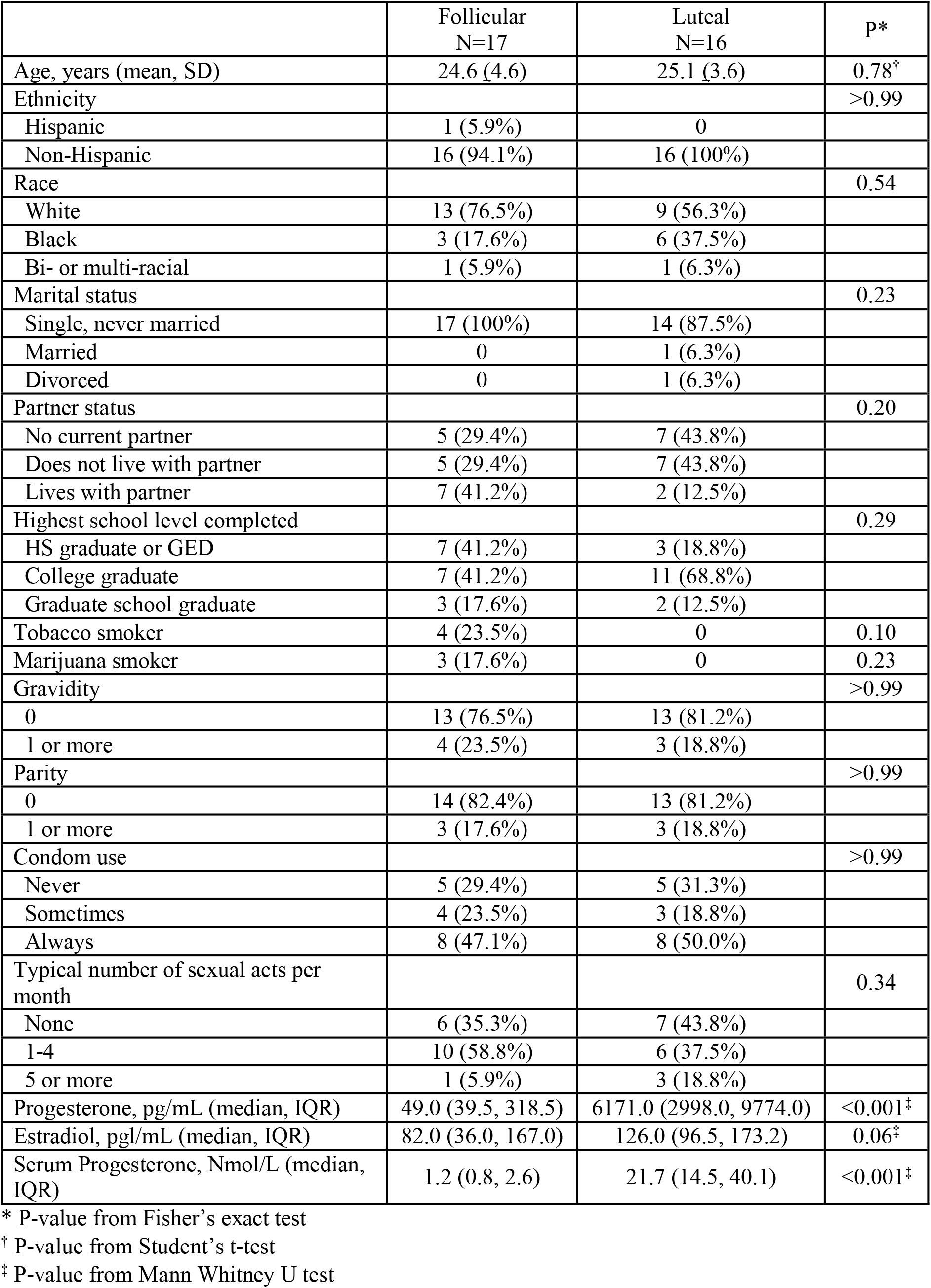
Demographic characteristics.

|  | Follicular<br>N=17 | Luteal<br>N=16 | P* |
| --- | --- | --- | --- |
| Age, years (mean, SD) | 24.6 (4.6) | 25.1 (3.6) | 0.78<sup>†</sup> |
| Ethnicity |  |  | >0.99 |
| Hispanic | 1 (5.9%) | 0 |  |
| Non-Hispanic | 16 (94.1%) | 16 (100%) |  |
| Race |  |  | 0.54 |
| White | 13 (76.5%) | 9 (56.3%) |  |
| Black | 3 (17.6%) | 6 (37.5%) |  |
| Bi- or multi-racial | 1 (5.9%) | 1 (6.3%) |  |
| Marital status |  |  | 0.23 |
| Single, never married | 17 (100%) | 14 (87.5%) |  |
| Married | 0 | 1 (6.3%) |  |
| Divorced | 0 | 1 (6.3%) |  |
| Partner status |  |  | 0.20 |
| No current partner | 5 (29.4%) | 7 (43.8%) |  |
| Does not live with partner | 5 (29.4%) | 7 (43.8%) |  |
| Lives with partner | 7 (41.2%) | 2 (12.5%) |  |
| Highest school level completed |  |  | 0.29 |
| HS graduate or GED | 7 (41.2%) | 3 (18.8%) |  |
| College graduate | 7 (41.2%) | 11 (68.8%) |  |
| Graduate school graduate | 3 (17.6%) | 2 (12.5%) |  |
| Tobacco smoker | 4 (23.5%) | 0 | 0.10 |
| Marijuana smoker | 3 (17.6%) | 0 | 0.23 |
| Gravidity |  |  | >0.99 |
| 0 | 13 (76.5%) | 13 (81.2%) |  |
| 1 or more | 4 (23.5%) | 3 (18.8%) |  |
| Parity |  |  | >0.99 |
| 0 | 14 (82.4%) | 13 (81.2%) |  |
| 1 or more | 3 (17.6%) | 3 (18.8%) |  |
| Condom use |  |  | >0.99 |
| Never | 5 (29.4%) | 5 (31.3%) |  |
| Sometimes | 4 (23.5%) | 3 (18.8%) |  |
| Always | 8 (47.1%) | 8 (50.0%) |  |
| Typical number of sexual acts per month |  |  | 0.34 |
| None | 6 (35.3%) | 7 (43.8%) |  |
| 1-4 | 10 (58.8%) | 6 (37.5%) |  |
| 5 or more | 1 (5.9%) | 3 (18.8%) |  |
| Progesterone, pg/mL (median, IQR) | 49.0 (39.5, 318.5) | 6171.0 (2998.0, 9774.0) | <0.001<sup>‡</sup> |
| Estradiol, pg/mL (median, IQR) | 82.0 (36.0, 167.0) | 126.0 (96.5, 173.2) | 0.06<sup>‡</sup> |
| Serum Progesterone, Nmole/L (median, IQR) | 1.2 (0.8, 2.6) | 21.7 (14.5, 40.1) | <0.001<sup>‡</sup> |

\* P-value from Fisher's exact test
<sup>†</sup> P-value from Student's t-test<sup>‡</sup> P-value from Mann Whitney U test

To characterise the spatial expression of IFNε in different parts of the FRT, we first analysed its expression using IHC in matched vaginal, ectocervical and endometrial biopsy samples from all participants. We found IFNε to be highly expressed in the stratified squamous epithelium localised to the basal and parabasal layers of the lower FRT (vagina and the ectocervix) (Figure 1A and B). In the endometrium, strong IFNε staining was detected in both the luminal and glandular epithelium (Figure 1C). In a stratified analysis based on menstrual cycle stage of participants, IFNε protein was significantly higher in the endometrium during the luteal phase of cycle compared with the follicular phase (Figure 2A, 2B). Conversely, there were no changes in IFNε protein expression across cycle stage in the vagina and ectocervix (Supplemental Figure 1).

**Figure 1.**
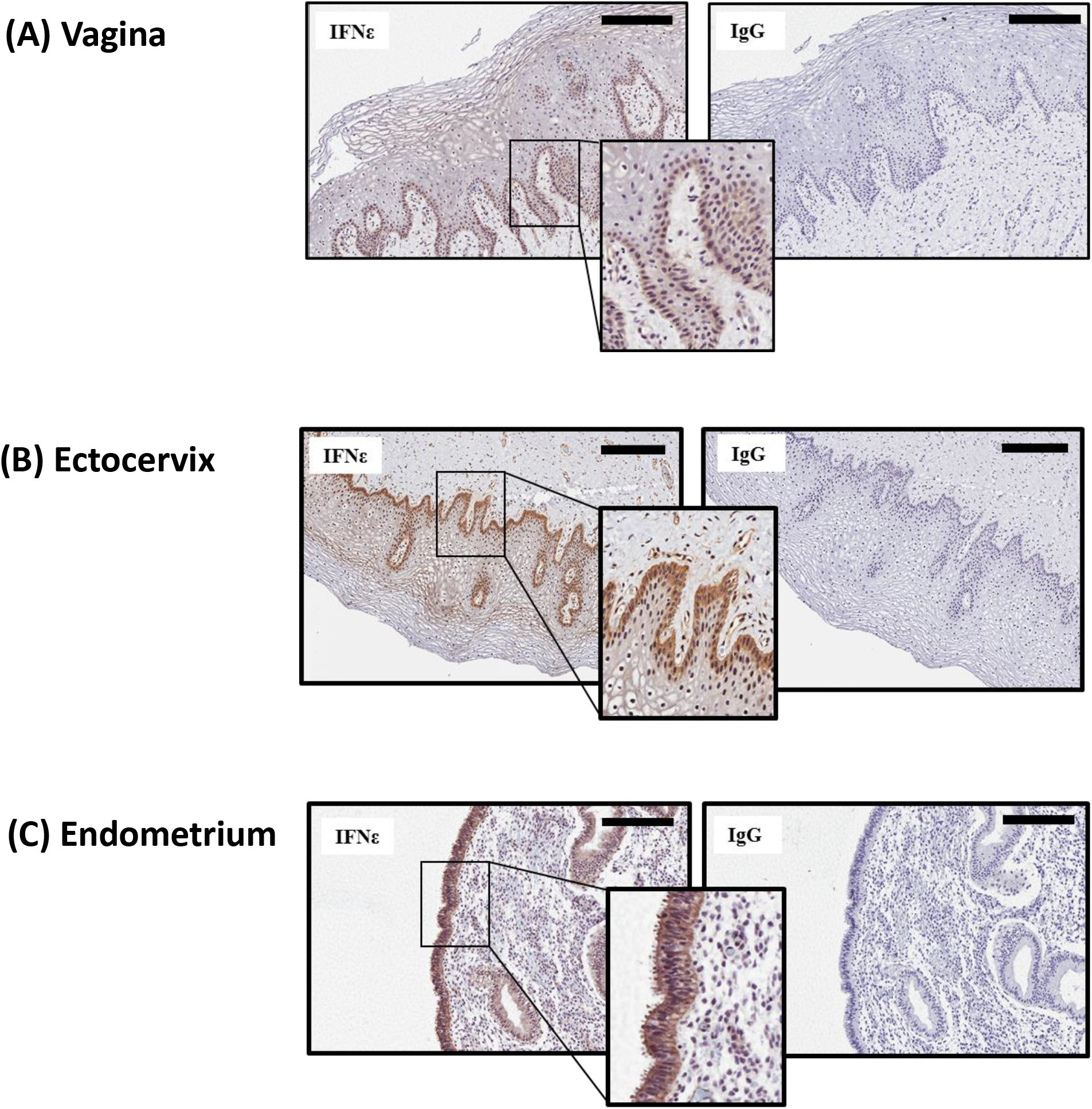
IFNε is expressed in distinct epithelial layers in the lower and upper human FRT. Representative images of IFNε expression in sections from matched biopsy samples from n=33 women. Sections from the (A) vagina (B) ectocervix and (C) endometrium were stained for expression of IFNε (brown) or IgG control. Bar, 200μM.

When *IFNE* mRNA was analysed in biopsy samples, there was significantly higher expression in luteal stage of cycle in the endometrium when compared with endometrial follicular stage expression (Figure 2C; p<0.01), a finding consistent with the protein expression data above. There was no change in vaginal and ectocervical *IFNE* expression between follicular and luteal stages of cycle. Thus, we established that IFNε is highly expressed throughout the human FRT including the basal layer of epithelium in the cervix and vagina. However, despite high constitutive expression of IFNε in the lower FRT, hormonal regulation was not observed in these locations, with IFNε only regulated in response to sex hormones within the endometrium.

**Figure 2.**
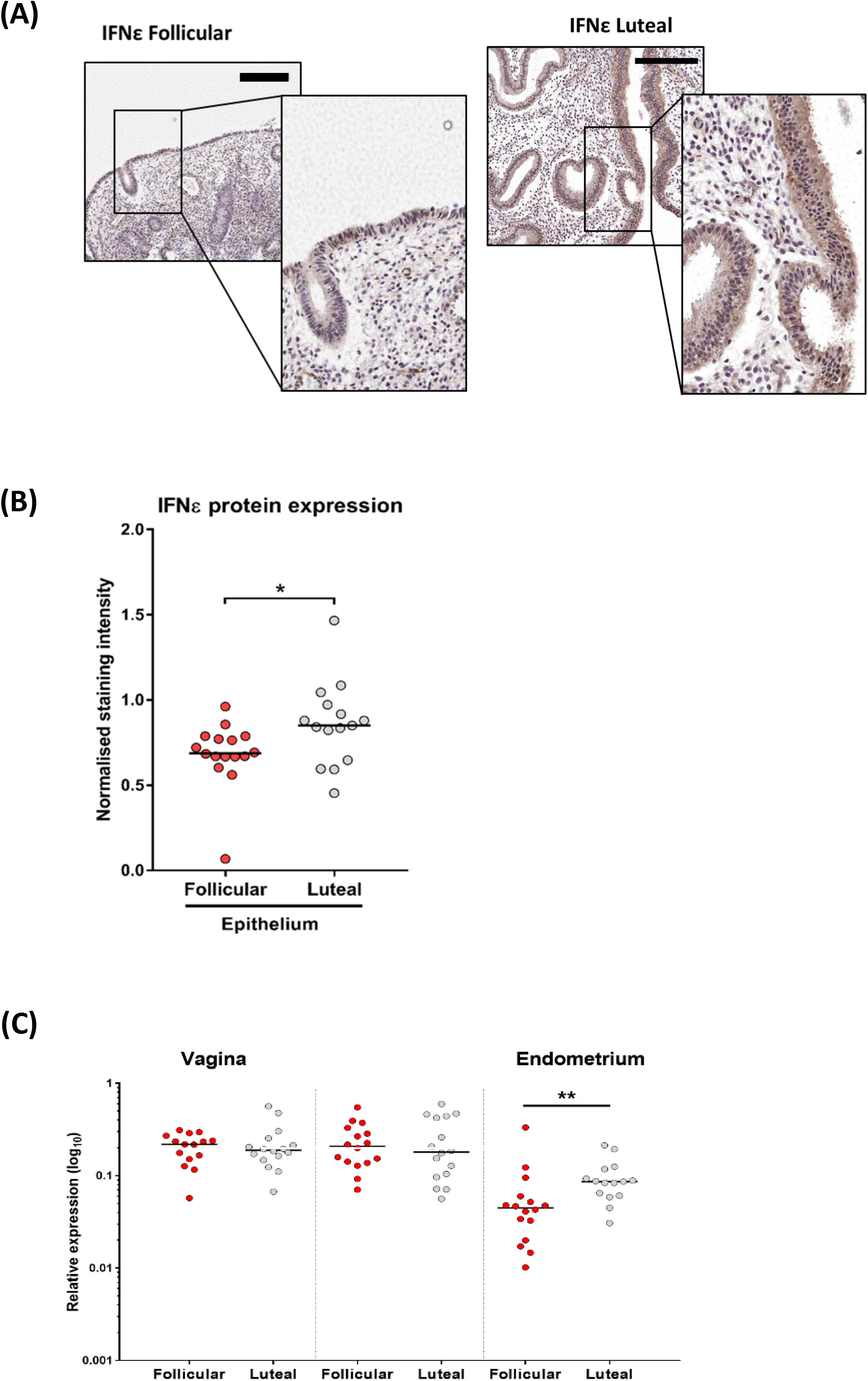
IFNε is hormonally regulated exclusively in the endometrium. (A) Representative IHC images of endometrial IFNε protein expression in follicular and luteal phases of menstrual cycle. Bar, 200μM. (B) Quantification of endometrial IFNε staining intensity in women in follicular or luteal stage of menstrual cycle. (C) *IFNε* mRNA expression, as determined by qPCR, in vaginal, ectocervial and endometrial biopsy samples stratified into follicular (n=16) and luteal (n=16) stages of menstrual cycle. Significance determined using Mann-Whitney U test; *p<0.05, **p<0.01.

### IFNε expression is negatively regulated by PR

Previous studies from our lab indicated that murine IFNε expression was negatively regulated by progesterone (Fung et al., 2013), but whether such regulation occurs in the human FRT was unknown. We found high expression of progesterone receptor (PR) protein (Figure 3A) and transcript (Figure 3B) in the endometrium relative to the very low to undetectable expression in the ectocervix and vagina. Further IHC analysis of PR expression across the FRT revealed strong staining of PR in both luminal and glandular endometrial epithelial cells. Furthermore, the significant reduction of PR expression in follicular stage of menstrual cycle seemed to particularly affect the cytoplasmic expression (Figure 3C and 3D). We also confirmed that the differential expression of PR with hormonal status in the endometrium was reflected in the mRNA levels, which reinforced the epithelial cell intrinsic nature of these differences (Figure 3E).

**Figure 3.**
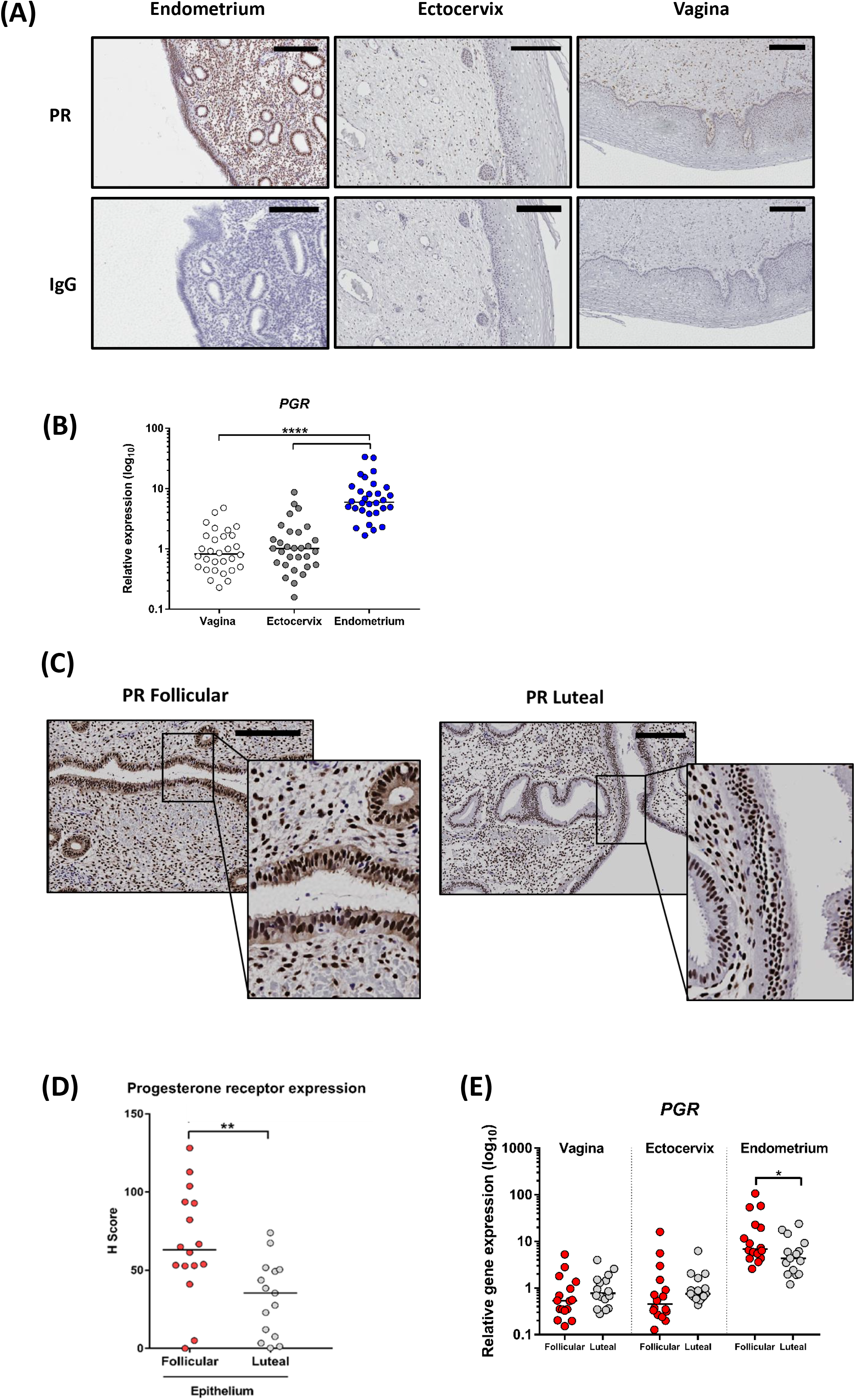
Progesterone receptor is hormonally regulated in the endometrium. (A) Representative images of PR expression in sections from matched biopsy samples from n=32 women. Sections from the endometrium, ectocervix and vagina were stained for expression of PR (brown) or IgG control. Bar, 200μM. (B) *PGR* mRNA expression, as determined by qPCR, in vaginal, ectocervial and endometrial biopsy samples stratified into follicular (n=16) and luteal (n=16) stages of menstrual cycle. (C) Representative IHC images of endometrial PR protein expression in follicular and luteal phases of menstrual cycle. Bar, 200μM. (D) Quantification of cytoplasmic PR staining intensity in endometrial epithelial and stromal cells from women in follicular or luteal stage of menstrual cycle. (E) *PR* mRNA expression, as determined by qPCR, in vaginal, ectocervial and endometrial biopsy samples stratified into follicular (n=16) and luteal (n=16) stages of menstrual cycle. Significance determined using Mann-Whitney U testing or Wilcoxon matched-pairs rank testing; *p<0.05, **p<0.01.

Since these independent measurements of IFNε and PR expression showed a pattern of inverse correlation, we directly characterised this apparent relationship in the cyclic changes in PR with IFNε expression in each sample and observed that IFNε expression in the FRT indeed showed a significant and inverse correlation with PR, a finding that was significant at both the mRNA and protein level (p<0.05; Figure 4A). Importantly, this effect was specific to PR expression as we found no correlation between *IFNE* and *ESR1*, the gene that encodes ERα, in the FRT, despite *ESR1* expression showing significantly lower levels in luteal phase of cycle (Supplemental Figure 2).

**Figure 4.**
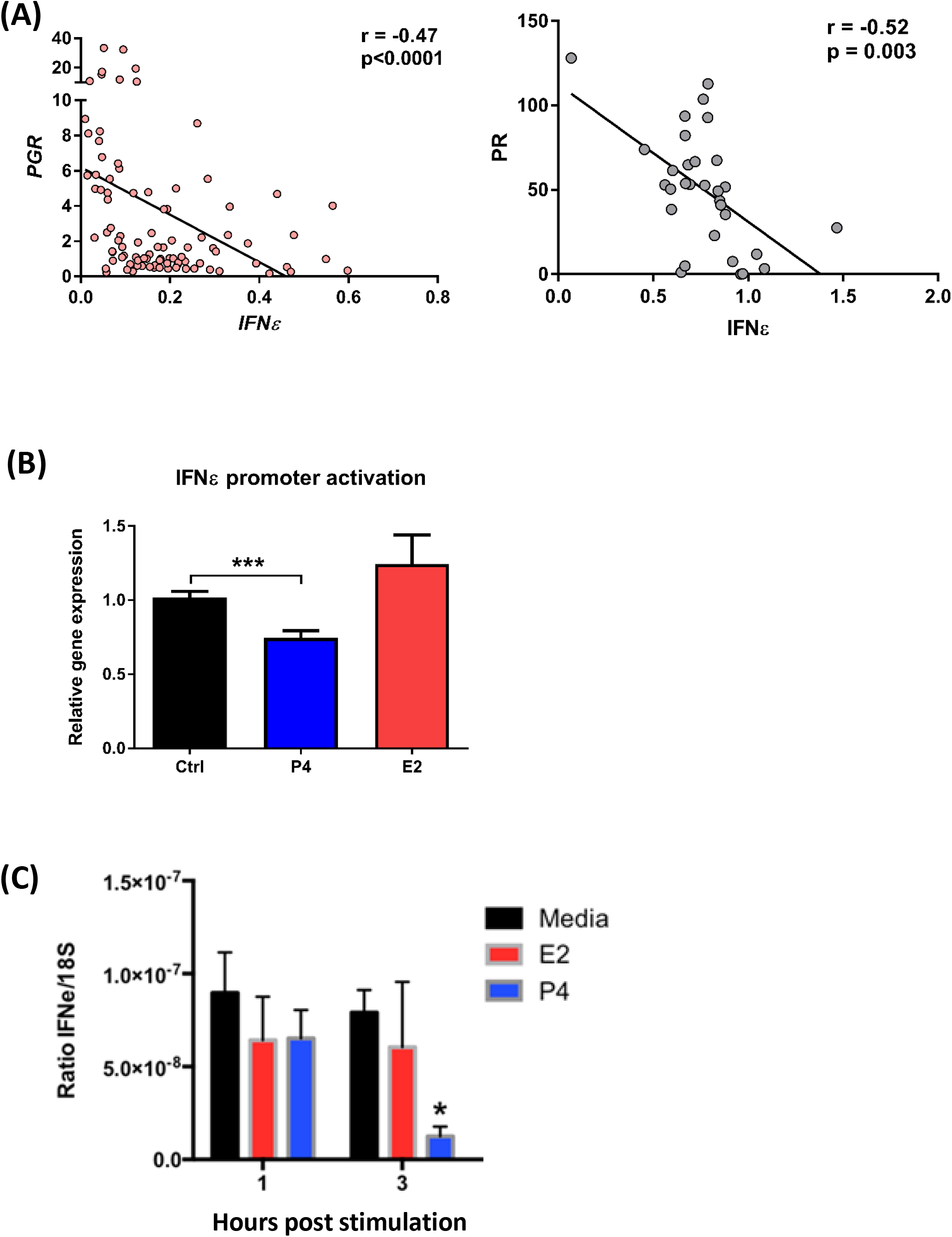
IFNε is negatively regulated by progesterone. (A) Negative correlation of both mRNA and protein expression of IFNε and PR in the FRT. Spearman correlation analysis. (B) Luciferase reporter assay measuring activation of the human IFNε promoter in ECC1 cells following treatment with 10^-9^ progesterone for 4h. Data from n=5 independent biological replicates, shown as mean +SD and analysed using Student’s T test, ***p<0.001. (C) Primary uterine epithelial cells were isolated from endometrial biopsies (from n=4 donors) and cultured for 3 days prior to stimulation with either 1 or 3 hours with 10^-9^ progesterone or 10^-9^ estrogen. *IFNE* expression was quantified using qPCR and expressed relative to expression of *18S.* Mann-Whitney U test, *p<0.05.

To directly investigate this implied inverse regulation of IFNε by PR, we characterised the regulation by progesterone of a luciferase gene construct under the control of the human IFNε promoter in an endometrial epithelial cell line, ECC1s. We found that stimulation with progesterone significantly inhibited expression of human IFNε (p<0.001; Figure 4C), yet estrogen stimulation had no effect on promoter activity. To further confirm this finding, we cultured primary uterine epithelial cells from four donor biopsies, stimulated them with progesterone and assessed *IFNε* expression. In this *ex vivo* model, *IFNε* expression significantly decreased following 3 hours of stimulation with progesterone (p<0.05; Figure 4D) whereas stimulation with estrogen did not alter IFNε expression. This data demonstrated the direct regulation of huIFNε expression *in vitro* by progesterone.

### IFNε regulates protective/immunoregulatory IRGs in the human FRT

We previously showed that IFNε protects against *in vivo* (HSV2 and Chlamydia) and *in vitro* (HIV) infection, which correlated with regulation of effector IRGs (Fung et al., 2013; Garcia-Minambres et al., 2017) We therefore sought to determine if huIFNε could regulate immune responses in the human FRT. We found that across the FRT, *IFNε* expression showed strong and significant correlation with expression of important immune IRGs, including *MX1*, *CXCL10*, *OAS2*, *IRF7*, *STAT1*, *DDX58* (Figure 5A). This thus fits with the hypothesis that basal expression of IFNε in the FRT constitutively maintains innate immune responses at this site. Although we found a positive correlation between IFNε and IRGs in FRT samples, there are many IFNs that can regulate these types of responses. However, the comprehensive and relative expression of all IFNs in the FRT had not been conclusively examined previously.

**Figure 5.**
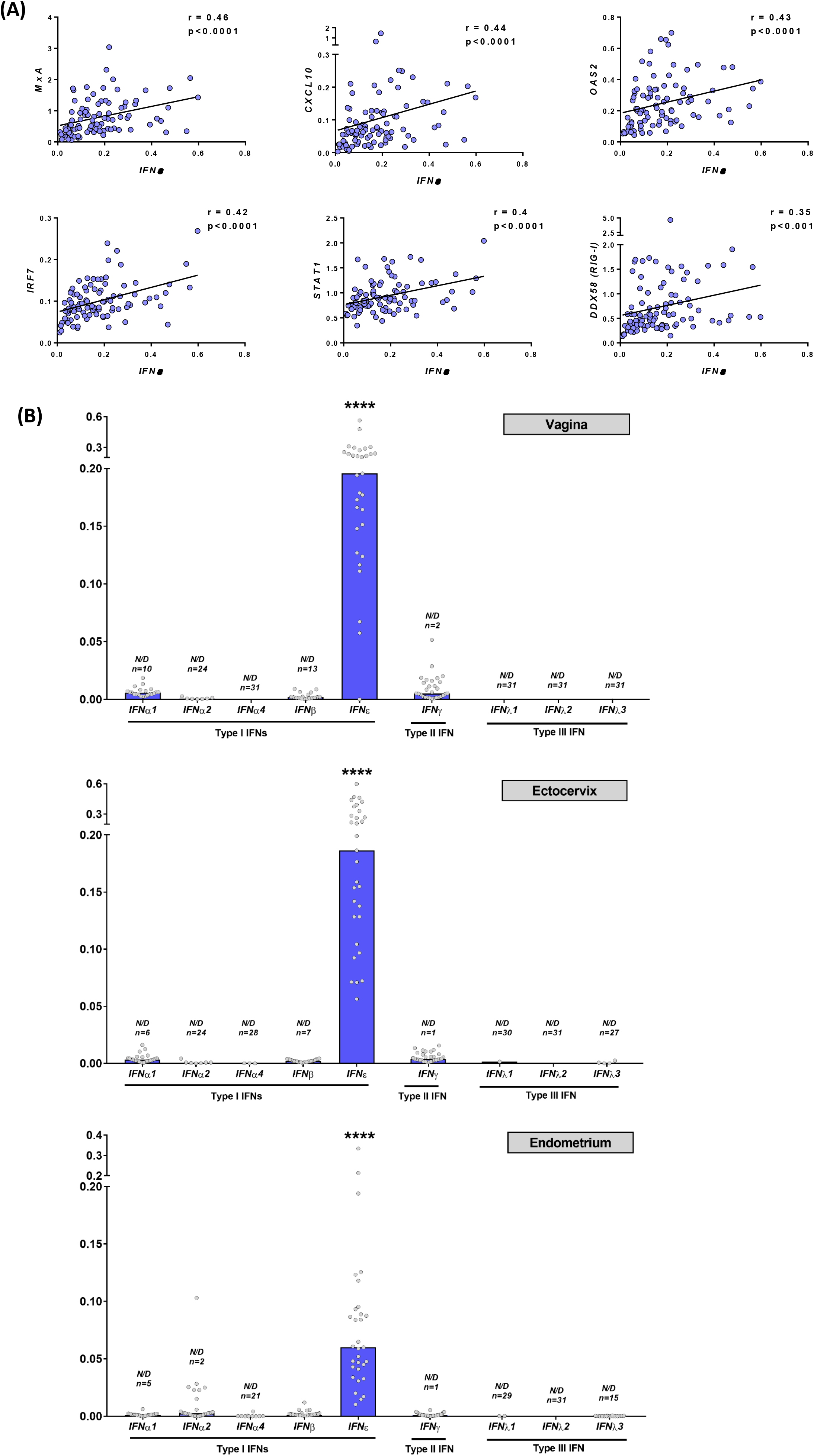
IFNε is the only IFN highly expressed in the human FRT. (A) Spearman correlation analysis of the expression of *IFNE* with the IRGs *MX1*, *CXCL10*, *OAS2*, *IRF7*, *STAT1* and *DDX58* across FRT samples. (B) Expression of type I IFN (*IFNA1*, *IFNA2*, *IFNA4*, *IFNB*, *IFNE*), type II IFN (*IFNG*) and type III IFN (*IL28A*, *IL28B*, *IL29*) was quantified by qPCR in paired vaginal, ectocervical and endometrial biopsy samples from n=32 women. N/D = not determined. Data analysed using Kruskal-Wallis testing with Dunn’s multiple comparison analysis, ****p<0.0001.

We therefore measured endogenous expression of other type I IFNs *IFNα1*, *IFNα2*, *IFNα4*, *IFNβ* and *IFNε*, the type II IFN *IFNγ* and the type III IFNs *IFNλ1*, *IFNλ2* and *IFNλ3* in matched biopsies by qPCR in all participants enrolled in this study (Figure 5B). Strikingly, we found that *IFNε* was strongly expressed in the vagina, ectocervix and endometrium when compared to the expression of other IFNs, a difference that was highly significant (P<0.0001). The expression of the other IFNs was mostly undetectable, except for low levels of IFNα1 and IFNγ in a few samples. Therefore, this data strongly suggests that IFNε is by far the key IFN maintaining IFN-dependent immunity in in the FRT.

We also assessed whether IFNε directly induced innate immune mediators in FRT epithelial cells *in vitro*. We stimulated vaginal epithelial cells (Vk2 cell line), ectocervical epithelial cells (Ect1 cell line) and uterine epithelial cells (primary cells cultured from endometrial biopsies) with IFNε and observed statistically significant induction of the IRGs *MX1, CXCL10* and *OAS2* (Figure 6A and 6B). Therefore, this data, along with the data in Figure 5, strongly suggests that IFNε is by far the key IFN maintaining IFN-dependent immunity particularly in homeostatic conditions in the FRT.

**Figure 6.**
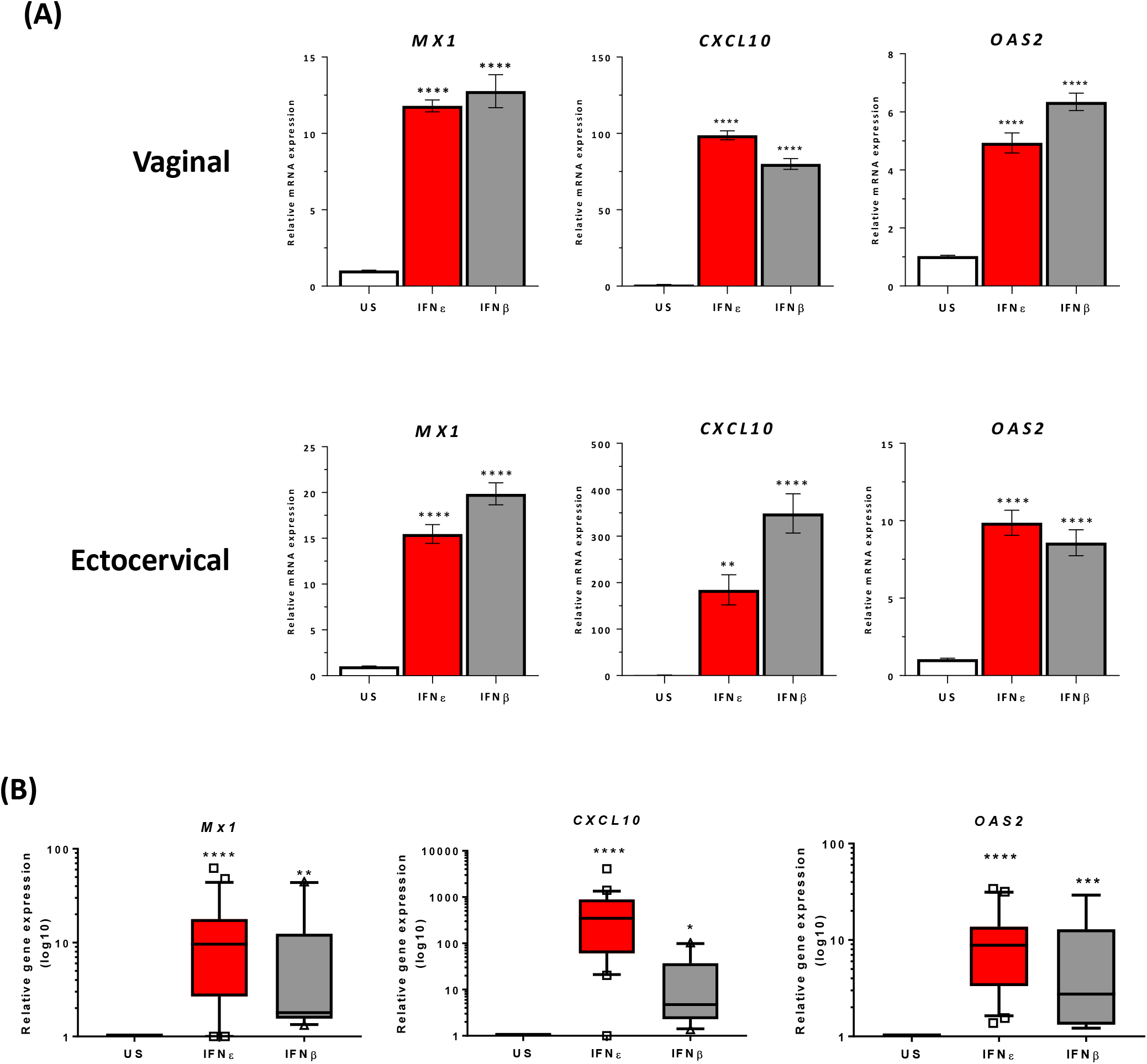
IFNε directly induces IRG expression in FRT epithelial cells. (A) Vaginal (VK2) and ectocervical (ECT1) cells were stimulated for 3hrs with 100 IU/ml of recombinant IFNε or IFNβ and expression of the IRGs *MX1*, *CXCL10* and *OAS2* were quantified by qPCR. Expression relative to unstimulated control. Data from n=3 independent biological replicates, shown as mean +SEM and analysed using Student’s T test using one-way ANOVA with Dunnett’s multiple comparisons testing **p<0.01, ****p<0.0001. (B) Primary uterine epithelial cells were isolated from endometrial biopsies and cultured for 3 days prior to stimulation with IFNβ (n=9) or IFNε (n=20). *IFNE* was quantified using qPCR, normalised to *18S* expression and expressed relative to untreated control cells. Mann-Whitney U Test, *p<0.05, **p<0.01, ***p<0.001, ****p<0.0001.

## Discussion

This study provides critical characterisation of the spatiotemporal expression and hormonal regulation of IFNε in distinct parts of the FRT in a cohort of 33 women forming a meticulously controlled and matched cohort. Importantly, the study design enabled definitive information in humans to complement our previous mechanistic studies in mouse models. We demonstrate that IFNε (mRNA and protein) is highly expressed in the luminal and glandular epithelium of the endometrium, the site of implantation and of immune importance for protection against ascending infections such as Chlamydia. There was even stronger expression of IFNε in the basal layers of the stratified squamous epithelium of the cervix and vagina, which are important sites of infection with viruses such as HIV and HPV. We and others recently demonstrated that IFNε can indeed directly inhibit replication of HIV *in vitro*. Therefore, there is continuous expression of IFNε from the lower to the upper human FRT at major sites requiring immune protection.

A clear finding in this study was that IFNε was the only IFN (type I, II or III) substantially and consistently expressed in all cases <u>constitutively</u> in the FRT. The absence of constitutive expression of type III IFNs is particularly important because the localisation of their receptor (IFNLR1) to epithelia cells has focussed a lot of attention on their role in protection of mucosal surfaces. At least here, in the uninfected state, IFNε is the sole IFN that will afford initial protection. The fact that immunoregulatory IRG expression in FRT samples was strongly correlated with the high IFNε expression present in the FRT indicates a compelling case for direct induction because IFNε was the only IFN constitutively expressed. Furthermore, we confirmed this direct relationship with complementary *in vitro* studies showing IFNε induction of these IRGs in cultured FRT epithelial cells form vagina, cervix and uterus. This data is in agreement with our previous data showing that IFNε^-/-^ mice have reduced IRG expression in their FRT (Fung et al., 2013). Maintaining ‘basal’ constitutive expression of regulatory IRGs which can modulate cellular processes such as metabolism, differentiation, proliferation, survival, angiogenesis etc. - in addition to their more prominent protective role in viral and bacterial infection and general immunoregulation – is apparently a role solely in the FRT for this unique IFNε.

This study defines that IFNε is hormonally regulated in the human FRT. Surprisingly, this regulation occurs exclusively in the endometrium and not in the vagina or ectocervix. This is consistent with our finding that progesterone responses directly suppress IFNε expression and that PR is also predominantly expressed in the upper FRT. Indeed, there is a substantial and significant inverse correlation between IFNε and PR expression at the mRNA and protein level across the 32 individuals in this study. While we didn’t observe hormonal regulation in the lower FRT, IFNε expression in the human ectocervix has been shown to be upregulated upon exposure to semen (Demers et al., 2014; Sharkey et al., 2007). This upregulation of ectocervical expression of IFNε in response to semen exposure, examined in a cohort of long term HIV-1 highly exposed, seronegative female sex workers, was hypothesised to be an immunomodulatory mechanism within the FRT to reduce risk of acquisition of HIV (Abdulhaqq et al., 2016). These results together with our data reported here, confirming constitutive IFNε expression not only in the ectocervix but throughout the FRT and that IFNε regulates IFN immunity in FRT epithelium, is consistent with IFNε likely representing an important basal FRT defence against pathogens.

Based on the evidence presented here, IFNε elicits IRG-driven pathophysiological actions in the FRT and therefore modulation of its expression in the FRT could have profound physiological consequences in terms of susceptibility to infection. For example, women on progestin-based contraceptives have been implied to be more susceptible to infection which may be due, at least in part, to reduce IFNε in the FRT.

## Acknowledgements

The authors would like to acknowledge Dr Rebecca Smith and Dr Deborah Bianco for assistance. The authors also acknowledge the scientific and technical assistance of Monash Histology Platform, Department of Anatomy and Developmental Biology, Monash University and the Monash Health Translational Precinct Medical Genomics Facility – Fluidigm Single Cell and High Throughput Centre. This work was supported by the Bill and Melinda Gates Foundation project OPP1108501.

## Supplemental figures

**Figure S1.**
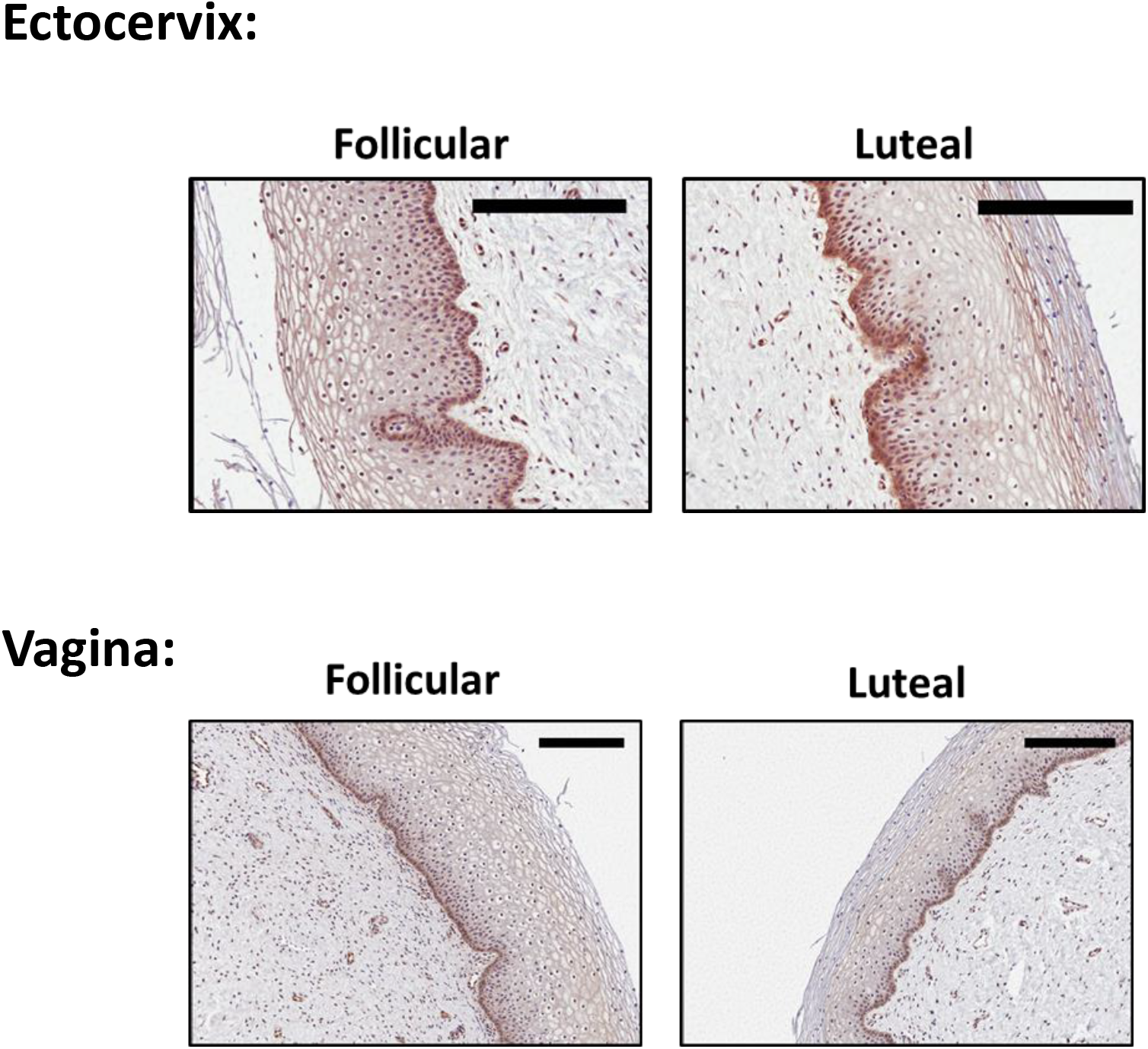
Staining of IFNε (brown) in vaginal and ectocervical sections in the follicular and luteal phase of menstrual cycle. Bar, 200μM.

**Figure S2.**
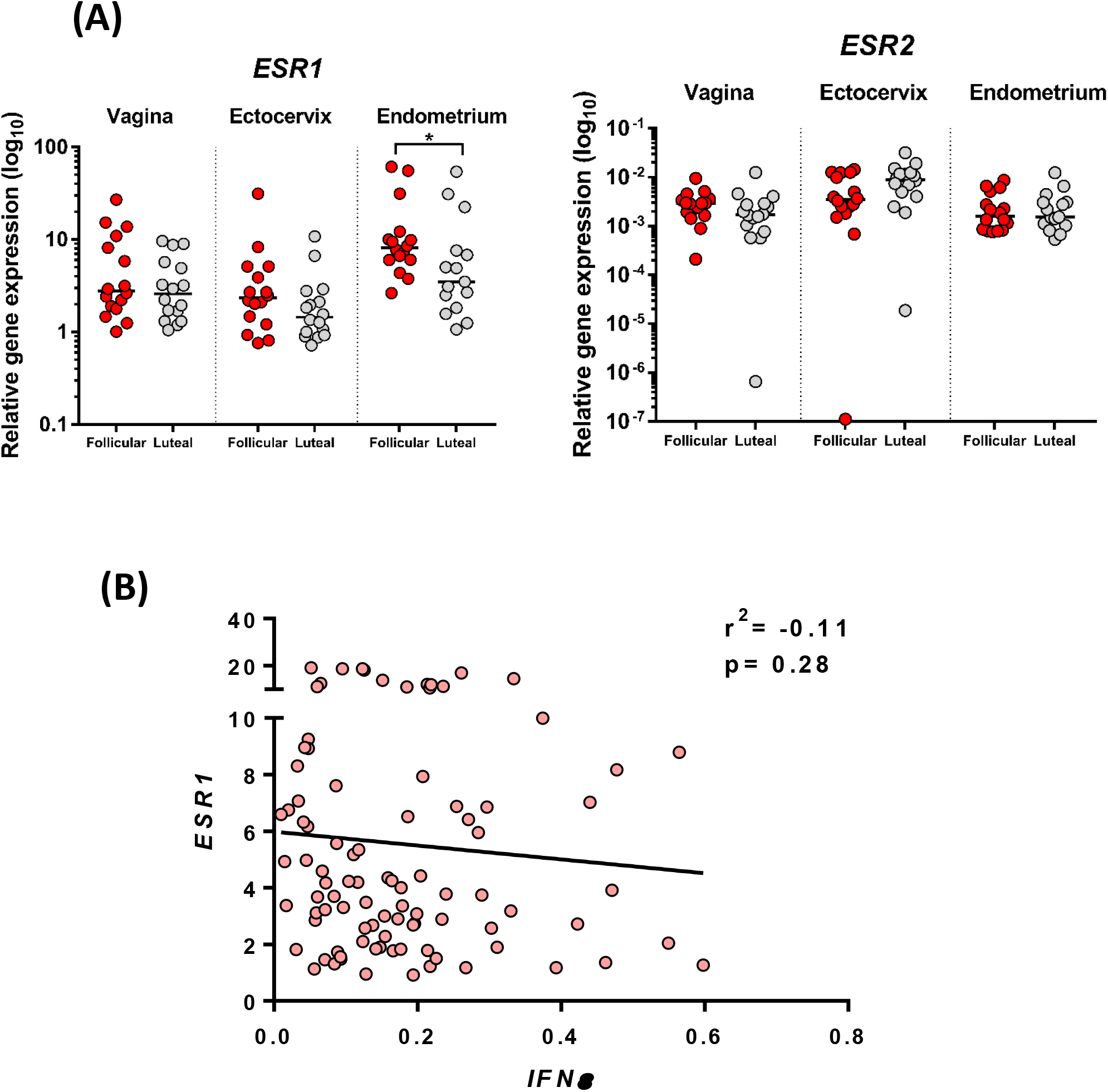
(A) Expression of *ESR1* and *ESR2* in vaginal, ectocervical and endometrial biopsies as determined using qPCR. Expression stratified by stage of menstrual cycle and median expression is shown. Data analysed using Mann-Whitney U-tests, *p<0.05. (B) Spearman correlation analysis of *ESR1* and *IFNE* expression in the FRT.

